# Lineage-associated Human Divergently-Paired Genes (DPGs) Exhibit Regulatory Characteristics and Evolutionary Trends

**DOI:** 10.1101/2024.12.10.625558

**Authors:** Guangya Duan, Sisi Zhang, Bixia Tang, Jingfa Xiao, Zhang Zhang, Peng Cui, Jun Yu, Wenming Zhao

## Abstract

Divergently-paired genes (DPGs) represent one of the minimal co-transcriptional units (the rest include tandemly- and convergently-paired genes) of clustered genes; the former and the latter constitute greater than 10% and 75% of the total human genes, respectively. Our previous studies have shown that vertebrate DPGs are more conserved, both organizationally and functionally than invertebrates. Three critical questions remain to be addressed: (1) what are the conserved DPGs over vertebrate lineages, especially among mammals and primates? (2) being bidirectionally transcribed, to what extent do DPGs share their promoter sequences and how mechanistically and stringently are their co-expression regulated within the shared inter-TSS (transcription start site) sequence space? and (3) based on the recently released high-quality human genome assemblies, how do human-associated DPGs distribute over selected primate lineages and what are their possible functional consequences biologically? Our study begins by identifying 1399 human DPGs (12% of all human protein-coding genes), and presents findings from this analysis. First, 1136, 1118, 925, and 830 human DPGs are shared genetically with primates, mammals, avians, and fish, respectively. DPGs are not only functionally enriched toward direct protein-DNA interactions and cell cycle synchronization but also exhibit obvious lineage association, narrow in principle toward synchronization of certain core molecular mechanisms and cellular processes. Second, their inter-TSS distances and expression variables affect both co-expression strength and disparity between the two genes. Finally, our results based on a comparison among the primate DPGs reveal that the human-associated DPGs exhibit intensive diversification in co-expression, duplication, and definite involvement in neural development. Within humans, 55 and 357 DPGs are associated to the Chinese (YAO) and the European (CHM13) assemblies, respectively. Our results offer novel insights into comprehending the structure-function selection of gene clusters over evolutionary time scales, as well as a deeper understanding of the regulatory characteristics of co-expressed neighboring genes.

## Introduction

In prokaryotes, gene clusters, commonly referred to as operons, represent a prevalent organizational formality where numerous non-randomly aggregated genes contribute to a singular function or phenotype [1,2]. Eukaryotes also exhibit a non-random distribution of certain gene families or classes, forming distinct structural configurations from primary sequences to three-dimensional topological structures of chromatins. Paired genes, as a form of minimal gene clusters, manifest in three configurations: divergently-, tandemly-, and convergently-paired [3,4]. Such genomic architectures are prevalent in diverse animal and plant lineages, underscoring their common manifestation [5–9]. It has been observed that over 10% of the human genes, i.e., divergently-paired-genes (DPGs), have a bidirectional orientation separated by less than 1000 bp [4,10]. Such a spacing constraint suggests advantages in sharing promoter elements and synchronized gene expression or co-expression of these DPGs, which may be functionally selected over an evolutionary time scale [6,11].

Comparative genomic analyses have indicated a high-degree of conservation and stability of this gene-pairing architecture among vertebrate genomes [5,12,13]. For instance, there are more fully-conserved DPGs among species of vertebrate lineages, when compared with those of the invertebrates as represented by insects [3]. Unlike the unidirectional model, DPGs are not formed from random chromosomal rearrangement events but show functional relevance across eukaryote genomes [14–16]. The formation of DPGs, by reorganizing existing genes, often leads to novel co-regulated transcription, representing a mechanism to acquire new components beneficial to species fitness [6,17,18]. Hence, the identification and study of lineage-associated (for association within or among lineages) and species-specific (for uniqueness in species) DPGs are essential in understanding diverse gene and genome evolution processes.

The detailed regulatory mechanisms of DPGs have recently been readdressed. The sequence-space-limited organization of DPG promoters places the two neighboring genes under pressure to share their regulatory elements at least in part, offering an opportunity to scrutinize how cis- and trans-elements interact in molecular details at the transcriptional level [19]. Several regulatory mechanisms, including epigenetic signals, have been identified, such as RNAPII occupancy and histone modifications of H3K4me2, H3K4me3, and H3ac, which are overrepresented at bidirectional promoters [20–22]. Additionally, the nucleosome-depleted regions (NDRs) between paired genes bring an optimal condition for co-regulation [11,23]. Furthermore, bidirectional promoters, commonly found among DPGs, often lack typical TATA boxes but possess high GC content and enrichment of CpG islands [4,24]. Moreover, they display distinctive sequence signatures characterized by a mirror composition, where Gs and Ts vs. Cs and As dominate the opposing sides [25]. The DPG-associated promoter sequences also influence transcription factor (TF) binding, and a small set of TF motifs appear to be overrepresented within the shared space [20].

As sequencing technology has advanced greatly in recent years, the quantity and quality of genomic data in various forms and from diverse sources have been increasing, allowing for in-depth studies, including orthologous relationships between genes, functional annotations, epigenetic modifications, RNA-seq data, and transcription factor motifs. Leveraging multiple-level high-quality datasets from diverse sources, we explore the evolutionary history and trajectory of DPGs across vertebrate lineages in a human-centric way and conduct a comprehensive comparative analysis based on primate data collections. In addition, taking advantage of the three recently-released human reference genomes, two of them are attributed to telomere-to-telomere (T2T) quality, i.e., T2T-CHM13 of European origin [26] and the T2T-YAO [27] of the first representative tailored to Chinese populations, we examine not only genomic disparities of the human DPGs across other vertebrate lineages and among human populations, but also analyze regulatory mechanisms of co-expressed DPGs, including genetic and epigenetic mechanisms, regulatory patterns of co-expression, shared TFs within inter-TSS sequences (referring to sequences in the defined space between two TSSs of DPGs), and other structural and functional features, to enhance our understanding of this important structural arrangement.

## Results

### Human DPGs exhibit complex lineage-associations

Utilized spacing-based criteria (inter-TSS between DPG < 1000 bp), we started our analysis with a set of 1,399 human protein-coding DPGs (Table S1). To investigate their evolutionary conservation patterns across several vertebrate lineages, we selected, aside from human, 45 species from 6 distinct lineages for further comparative analysis (Figure 1 and Table S2). Four conservation categories were defined to quantify the DPG conservation levels (see Materials and Methods for details). Among the lineages and species, the four conservation categories are in similar proportions, varying based on sequence identities relative to the human genes, but their precise numbers may be difficult to define due to the quality of their genome sequences. For instance, gorillas and chimpanzees show the highest numbers of highly conserved human DPGs, which is to be expected, but appear to show fewer conserved DPGs when compared to those of the mouse due to the better murine genome assembly. Three invertebrate species, *Drosophila melanogaster, Caenorhabditis elegans*, and *Saccharomyces cerevisiae* were also included in our analysis as representative outgroups (Figure 1A). A hierarchical clustering mapped the species based on their sequence-based similarity distance in each row and displays them as three major groups: the mammals stand-alone, fishes and amphibians are grouped together, and avians form the third group. The reptiles split into either avians or the lower vertebrate groups, and such a deviation from expectation is attributable to two reasons at least; one, the small number of reptilian species used for the analysis, and the other, the number of DPGs are species- and lineage-associated, which are partly convergently evolved and selected for their functional significance (Figure 1B).

**Figure 1.**
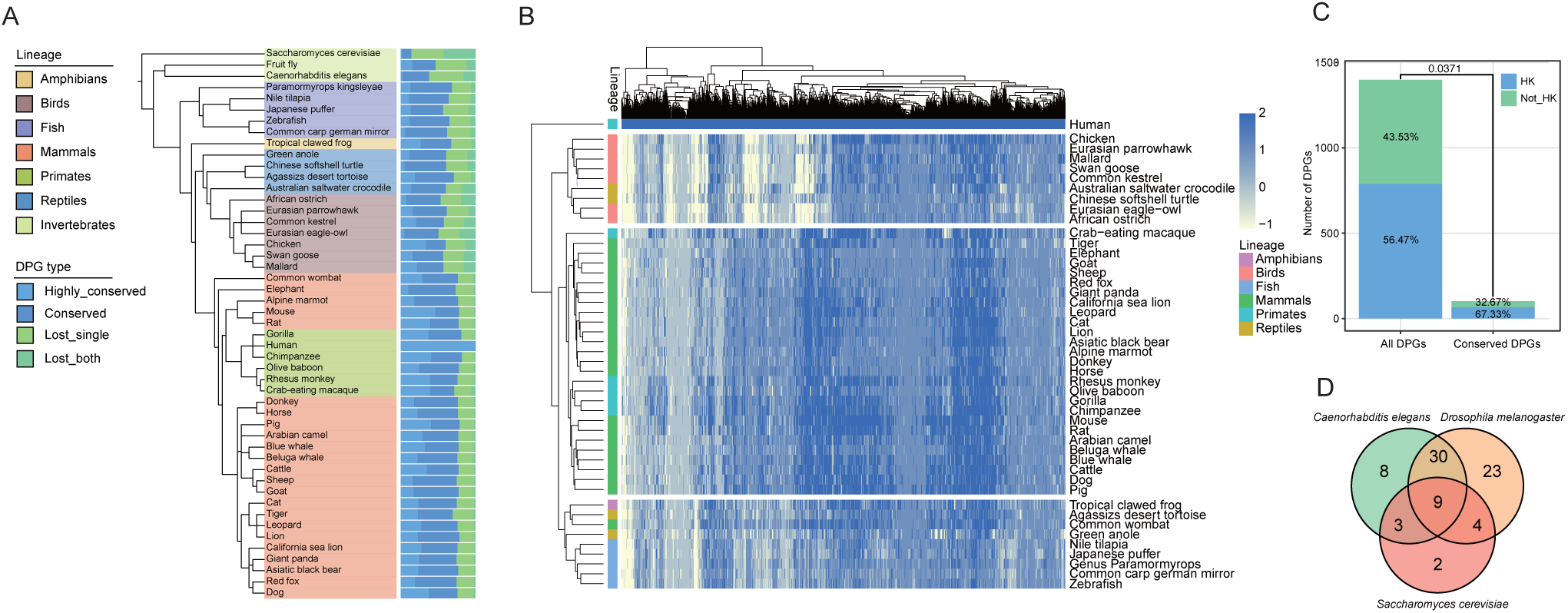
The four conservation forms of human DPGs across diverse vertebrate lineages. **A.** Fractions of the four gene- and orientation-defined forms of conserved DPGs from different species. The organization is based on phylogenetic tree originated from NCBI taxonomy. The detailed categories method is shown in Materials and Methods. **B**. A heatmap showing the conservation of human DPGs in 45 other vertebrates, where darker colors indicate higher conservation. The values of 2, 1, 0, and −1 correspond to the four conservation types, representing a conservation scale from high to low. The columns and rows are grouped by two-way hierarchical clustering. **C.** The proportion of DPGs containing house-keeping (HK) genes of all human protein-coding DPGs and in the 101 vertebrate-conserved DPGs (vcDPGs). **D.** A Venn diagram illustrating the conservation of vcDPGs in three selected eukaryotic multi-and unicellular species, *Drosophila melanogaster, Caenorhabditis elegans*, and *Saccharomyces cerevisia*.

Conversely, most lineage-associated DPGs appear to follow a divergent trend, where the 1399 human protein-coding DPGs share 1136, 1118, 830, and 925 with other primates, mammals, birds, and fishes, respectively. We even have 101 vertebrate-conserved (vcDPGs) over all 46 selected species for this study.

We also looked into two other interesting aspects, partitioning between house-keeping (HK) and tissue-specific (TS) functions and conservation across distinct lineages. Using a list of HK genes as benchmarks from a previous study [28], we found that DPGs are slightly HK-gene enriched, harboring 56.47% of all and 67.33% of vertebrate-conserved DPGs (Figure 1C), whereas the general partition of HK and TS genes are half-and-half [29]. There are 67, 50, 18, and 9 conserved in the fruit fly, the nematode, the yeast, and all three species, respectively, showing conservation of these gene pairs even among invertebrates (Figure 1D, Table S3).

### Functional relevance of vcDPGs

The functional relevance of DPGs has long been known as they must follow lineage-associated characteristics defined based on evolutionary principles and criteria [30,31]. Since DPGs stand out as a unique class of co-expressed genes for genome-wide functional and mechanistic analyses, we decided to take a subgroup of them as an example for in-depth exploitation, and thus the choice of vcDPGs. Our simple enrichment analysis suggests “binding” functions in the Molecular Function category, such as protein-RNA binding. In the category of Biological Process, the enrichment appears to relate to DNA repair, cell cycle, and intracellular protein transport. As to the Cellular Component category, what is enriched includes the nucleus, cytoplasm, cytosol, and nucleoplasm (Figure 2A). These results suggest that vcDPGs, belonging to the most conserved group of genes over 500 million years of vertebrate evolution [32,33], are most likely to be associated with certain fundamental binding functions that coordinate or synchronize the timing of cell-cycle-related processes through direct interactions among proteins, RNAs, and DNA. In other words, these proteins tend to play regulatory roles instead of evolving into functional proteins enriched in particular pathways.

**Figure 2.**
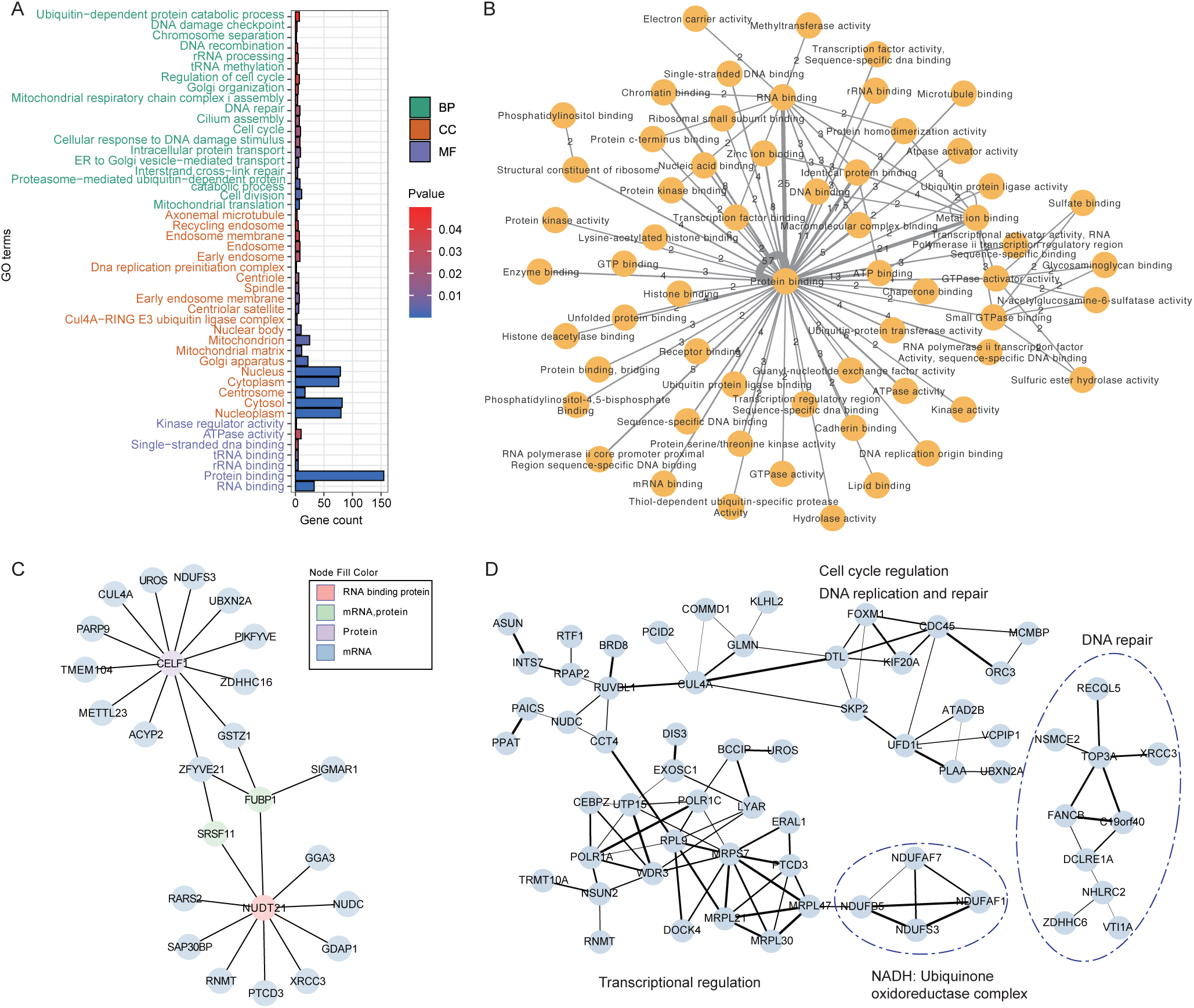
Functional interactions of vcDPGs. **A.** Functional enrichment of Gene Ontology annotations of the 101 vcDPGs obtained from DAVID. The bar color is filled based on P values, and the y-axis is colored based on the category. BP, biological processes, MF, molecular functions, CC, cellular components. **B.** Functional correlation network of vcDPGs, filtered with a minimum correlation count greater than 1. Each node represents a Gene Ontology (GO) term of the MF category, while each edge denotes a functional correlation between vcDPGs. Circles indicate vcDPGs with both genes sharing the same GO term. The numbers on edges indicate DPG count associated with that correlation. Most vcDPGs have functional correlations with the GO term “protein-binding”, as indicated by the thick circle. **C.** Protein-RNA interaction network of vcDPGs, filtered with a high confidence threshold using data obtained from the RNAInter database. The nature of the molecules is color-coded. **D.** A PPI network of vcDPGs. The data are filtered based on a medium confidence threshold, preserving the primary clusters. All interactions are listed in Table S4. The clusters are composed of two major groups, one on the left and a separate group on the right.

To provide further insight, we investigated the Molecular Function category based on functional networking that highlights 57, 21, 21, and 11 binding details as protein binding, protein-RNA binding, protein-metal ion binding, and 11 protein-DNA binding, respectively (Figure 2B). From this list of genes, we learned that these bindings are not only physical interactions between or among certain proteins but are mostly involved in catalytic activities, such as phosphorylation and other post-translational modifications (PTMs). We constructed both RNA-protein interaction and protein-protein interaction (PPI) networks for further illustration. For instance, our RNA-protein interaction network analysis reveals that *CELF1* (CUGBP Elav-like family member 1) and *NUDT21* (Nudix hydrolase 21), both functioning as RNA-binding proteins, are involved in non-coding RNA and mRNA processing, respectively, and partitioned into two distinct yet related networking clusters (Figure 2C). Surprisingly, the counterparts that form DPGs with these two genes are *PTPMT1* (protein tyrosine phosphatase, mitochondrial 1) and *OGFOD1* (2-oxoglutarate and iron-dependent oxygenase domain containing 1), respectively, which are enzymes of different cellular functions. Similarly, most DPGs with protein-ion interactions are predominantly involved in zinc binding, with a few exceptions, mostly interacting with bivalent ions, such as calcium, copper, and magnesium ions (Table S4). These highly conserved vcDPGs appear diverse in functions but may orchestrate protein-prosthetic complex formation yet essential or cellular housekeeping processes. The final examples are composed of two PPI clusters; both are related to DNA repair and also linked with RNAPII, mitochondrial ribosomal proteins, and subunits of ubiquinone oxidoreductase (Figure 2D, Table S4). To investigate direct interactions between the gene pairs among these DPGs, we compared the collection to 101 randomly selected gene pairs. In such a sampling test, there is only 1 out of the 101 randomly selected pairs exhibited direct protein interactions, whereas 41 out of the vcDPG pairs showed such an interaction. This result suggests that both the spatial proximity of DPGs and their coordinated expression regulation contribute to their functions, mostly selected for direct molecular interactions (P = 1.347E-13, Chi-squared test).

### Human DPGs show distinct tissue specificity and co-expression patterns

Although co-expression of DPGs has been extensively observed [31,34], their tissue specificity and the degree of such a coordinated cellular activity need further examination at a single DPG level. We conducted co-expression analysis for all 1399 DPGs across 51 tissues (Figure 3A). Our first impression is that DPG expression in brain-related tissues manifests higher co-expression coefficients as expected for TS genes, and some DPGs show opposing expression trends in different tissues. For example, expression of the *GSKIP-ATG2B* pair shows a positive correlation of 0.87 in brain-putamen (basal ganglia) and heart-left ventricle tissues, but a negative correlation of −0.6 in colon-transverse and vagina tissues becomes evident (Figure S1A). Conversely, the colon has the fewest co-expressed DPGs, totaling only 6. Compared with 1,399 randomly selected gene pairs from the human genome, DPGs exhibit a significant co-expression effect, particularly in brain-related tissues (P = 7.89E-121 and P = 1.267E-2, Chi-Squared Test, Figure 3B). After removing DPGs that are HK genes, the result becomes more pronounced (P = 2.778E-4, Chi-Squared Test, Figure 3C), highlighting the fact that tissue specificity and co-expression are both essential characteristics of DPGs.

**Figure 3.**
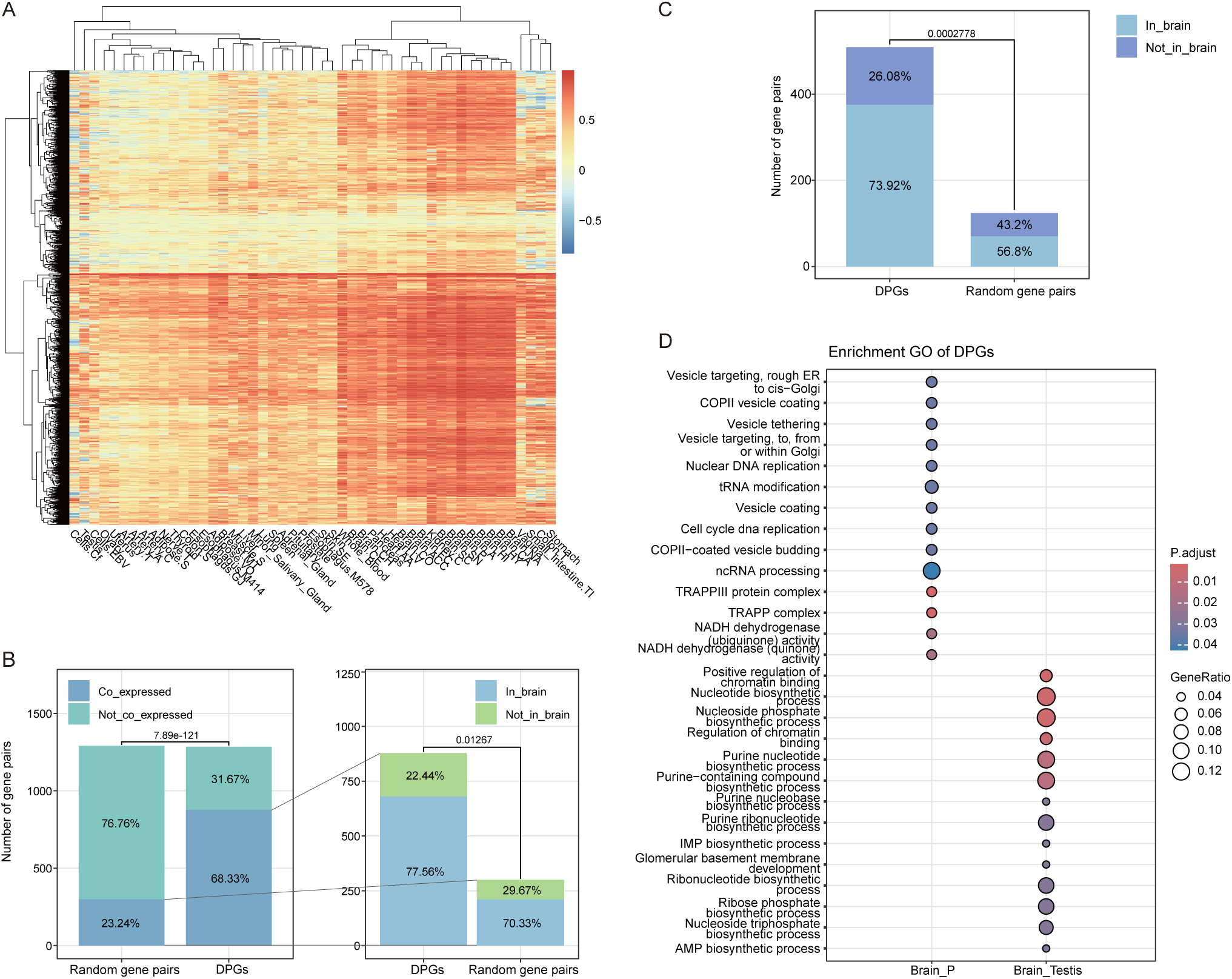
Tissue specificity and co-expression of DPGs. **A.** Correlation coefficient between expression values of DPGs across all tissues. The rows of DPGs and columns of tissues are grouped by two-way hierarchical clustering. Simplified names represent the tissues, and the full names are detailed in Table S6. **B.** The number and proportion of co-expressed DPGs in comparison to randomly selected gene pairs. Zoom in to the co-expressed proportions, the number and proportion of DPGs that co-expressed in brain-related tissues compared to randomly selected gene pairs. **C.** Comparison after removing HK genes. Chi-Squared Test was used to compute the difference in B and C. **D.** Functional enrichment of DPGs co-expressed in Brain-P tissue and DPGs co-expressed in both brain and testis tissues.

A total of 14 co-expressed DPGs are ubiquitously found in all 51 tissues, such as the *DTX3L-PARP9* and *TMEM176A-TMEM176B* pairs. These gene pairs are functionally enriched in essential biological processes, including structural integrity (e.g., basement membranes, collagen), protein homeostasis (e.g., proteasome function) and immune response regulation (e.g., dendritic cells, virus defense, BH adjusted P < 0.05, Figure S1B, Table S5). Others are tissue-specific, co-expressed in brain-putamen (basal ganglia) tissues and enriched in ncRNA processing, mitochondrial functions, and COPII vesicle coating. Interestingly, a subset of DPGs is co-expressed in both testis and brain tissues, enriched in nucleotide biosynthetic processes (BH adjusted P < 0.05, Figure 3D). These results suggest that tissue specificity and co-expression of lineage-associated DPGs should be considered separately with special attention.

### Highly co-expressed DPGs entertain inclusive regulatory mechanisms

To provide high-resolution views on regulatory mechanisms for DPGs, we incorporated four distinct epigenomic signal types, DNase, RNAPII, H3K4me3, and H3K27ac, along with sequence features such as GC count, using a sliding window approach (Figure 4A). High levels of RNAPII and DNase signals between the divergently transcribed genes of a DPG indicate increased chromatin accessibility, whereas H3K4me3 and H3K27ac signals highlight actively transcribed regions. These epigenetic marks reveal a regulatory landscape within the shared intergenic sequence of DPGs.

**Figure 4.**
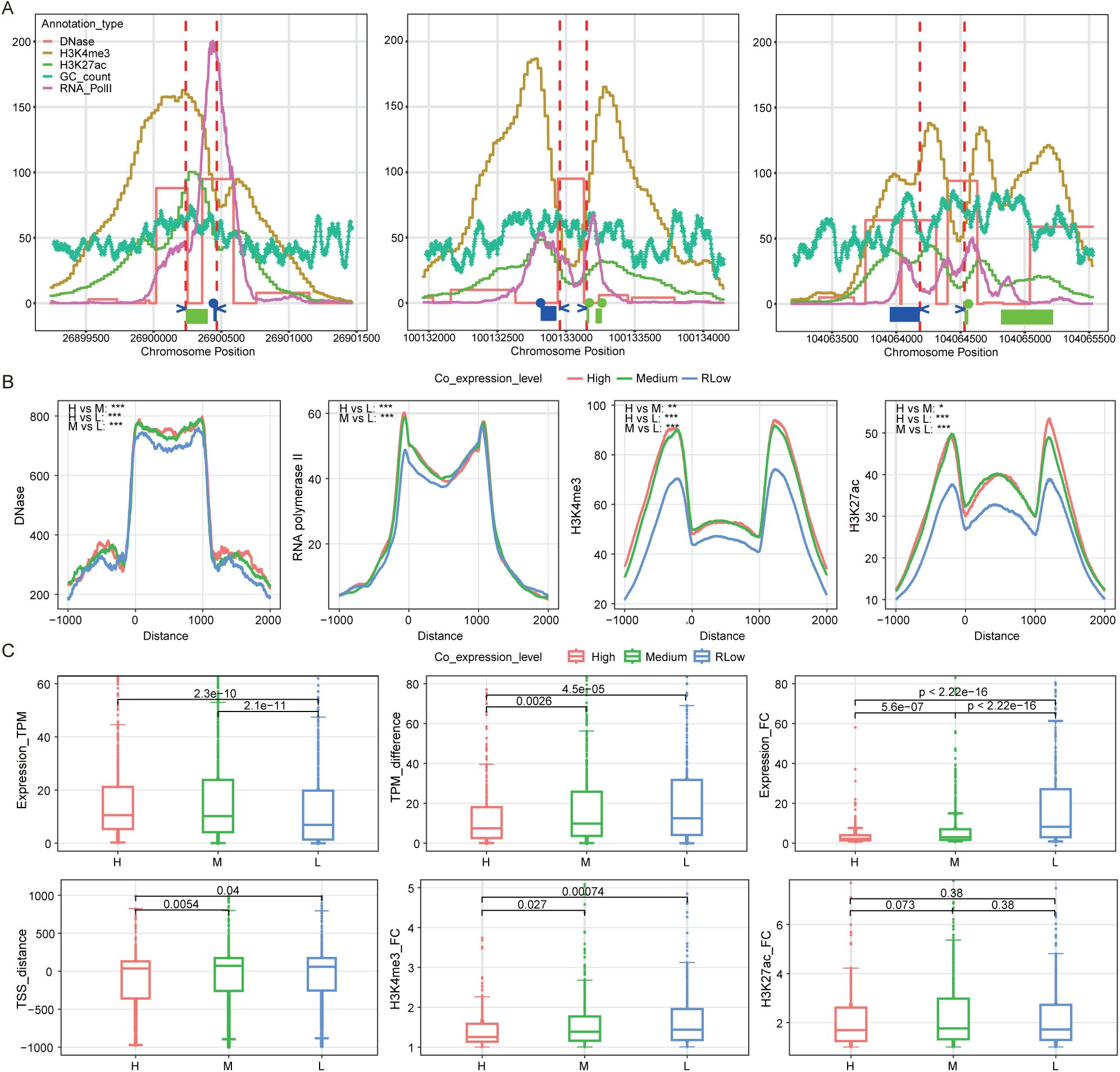
Epigenetic regulatory features of co-expressed DPGs. **A.** An illustration of diverse regulatory signals in DPGs. Regulatory signals are color-coded. The two vertical red dashed lines depict TSSs of the two genes; the blue arrows indicate transcription directions. The green horizontal line represents the 5’-UTR of genes on the positive strand, and the blue line represents the 5’-UTR of genes on the negative strand. Colored dots signify the start site of the first CD (coding DNA). **B.** Average curves illustrating the DNase, RNAPII, H3K4me3, and H3K27ac signals of DPGs at different co-expression levels. The curve distributions, predominantly corresponding to the groups ranging from High to RLow, mostly display positive correlations. **C.** Box plots illustrating characteristics of DPGs at different co-expression levels, including expression values, expression value differences, fold changes in expression, distances between TSSs, and fold changes in H3K4me3 and H3K27ac signal intensities. The first five figures only show P values that indicate significant between-group differences. The final figure of H3K27ac shows no inter-group differences. Statistically significant differences are denoted as follows: *P < 0.05, **P < 0.01, ***P < 0.001.Wilcoxon-Mann-Whitney U test was used to compute the difference in both B and C.

To find unique regularity characteristic of highly co-expressed DPGs, we grouped all DPGs into three levels based on co-expression correlation coefficients ranging from high to relatively low (RLow) and obtained mean curves for each epigenetic signal within each group. We observed positive correlations, in most cases, between co-expression levels and values of the four epigenetic signals (P < 0.05, Wilcoxon-Mann-Whitney U test, Figure 4B). As these signals signify active transcription, we further analyzed the expression disparity between the groups. Our results demonstrate that highly co-expressed DPGs exhibit a distinct set of characteristics. First, when compared to other DPGs, they show higher enrichment of DNase and RNAPII signals, as well as histone modifications, such as H3K4me3 and H3K27ac. Second, the genes forming the pair not only exhibit higher expression levels but their levels are rather comparable. Lastly, they have shorter or more overlapped inter-TSS distances as compared to those of the other two groups (P < 0.05, Wilcoxon-Mann-Whitney U test, Figure 4C).

### Sequence spacing influences DPG co-expression modes

H3K4me3 modification signals are markers for active promoters [35]. We further used peak positions to explore their regional distributions in DPGs. Overall, three distinct patterns were identified; each corresponds to unique inter-TSS spacing distributions: overlapping, optimal, and distant (Figure 5A and B). The numbers of DPGs in the three groups were 390, 817, and 149, respectively. The distant group constitutes the smallest proportion among all DPGs, particularly diminishing in highly co-expressed DPGs (Figure 5C), indicating a disadvantage in co-expression when the paired genes are too far apart from each other. In contrast, the overlapping and optimal groups of highly co-expressed DPGs are higher in proportion. The overlapping DPGs appear to be highly expressed individually and strongly co-expressed albeit with greater differences in expression levels among the pairs (Figure 5D and E). This suggests that promoter space competition may occur when the two genes are highly overlapped or crowded spacially.

**Figure 5.**
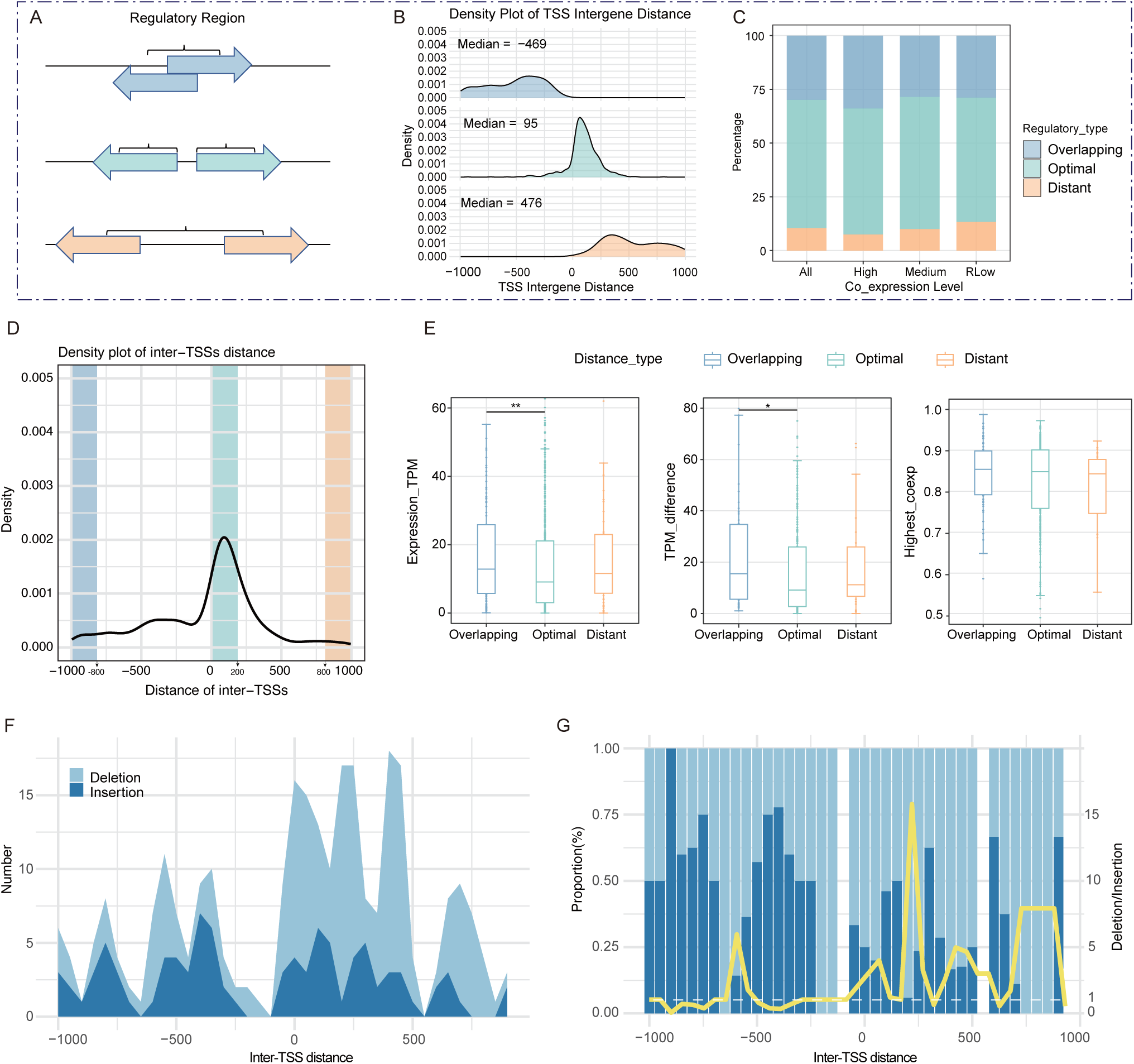
Sequence spacing within the regultory regions of DPGs affects sequence stability and co-expression. **A.** An illustration for inter-TSS distance or regulatory sequence spacing of DPGs. The spacing features are qualitatively classified into three basic types: overlapping, optimal, and distant. **B.** A diagram illustrating inter-TSS distances varied among DPGs in the three groups**. C.** The proportion of three types of DPGs in different co-expression level groups. **D.** The relative abundance of DPGs in the three types. **E.** Box plots illustrating expression values, differences in expression values, and co-expression of the three DPG types (the groups for comparisons are highlighted in D). Wilcoxon-Mann-Whitney U test was used to compute the difference. **F.** The number of filtered indels (see Materials and Methods for more details) plotted against in the inter TSS spacing. DPGs are partitioned based on inter-TSS distance with each data bin composed of 50-bp sequences. **G.** The proportion of insertions and deletions in each bin. The yellow curve shows the deletion/insertion ratio, and the number of insertions is set to greater than 1 for calculation.

Therefore, together with results from epigenetic signals and expression variables, our analyses reveal that the regulatory pattern of the co-expressed DPGs follows three distance-sensitive modes. First, when inter-TSS distances are appropriately situated within a 95-bp window in median, the majority of these DPGs, i.e., the optimal group, exhibit less biased co-expression and lower expression levels, and both are attributable to the shared open chromatin status and enriched cis-regulatory elements. Second, when inter-TSS distances are highly overlapping, more than a quarter of these DPGs (a median of −469 bp and extending to −1000 bp), the overlapping DPGs appear to be highly expressed individually and stronger co-expressed albeit greater differences in expression levels among the pairs. As inter-TSS distances increase (a median of 976 bp and extending to 1000 bp), the distant DPGs become apart from each other and their expression becomes less synchronized (P < 0.05, Wilcoxon-Mann-Whitney U test, Figure 5E).

To confirm genetic stability of the three DPG grouping schemes, we analyzed insertions and deletions (indels) in the inter-TSS sequence spacing across different DPG groups, using data from the expanded 1000 Genomes Project [36]. When the indels are plotted in a 50-bp sliding widow, the selective trends are clearly demonstrated (Figure 5F and G). There are relatively more deletions in the optimal and distant DPGs, indicating deletions contribute more to the overall indels in these two types; the opposite trend is observed for the overlapping group, where higher proportions of insertions are observed. Additionally, the distribution of insertion events appears relatively uniform other than the obvious peak-and-valley patterns that are attributable to nucleosome positioning. Taken together, these results suggest that inter-TSS spacing between DPGs influences sequence stability and gene expression variability. Deviations in TSS spacing within DPGs lead to more sequence-space-altering or sequence-length-sensitive mutations.

With no major differences in the numbers or types of TF-binding sites among the three groups, we identified two clusters of transcription factors (TFs) that are commonly shared among DPGs (Figure S2A). One functions in cell cycle monitoring, including G1/S phase transition and transcription activity, and the other involves epigenetic regulation of gene expression (Figure S2B and C). Taking vcDPGs as examples, by selecting the three most co-expressed TFs for each DPG as the core TFs, we show a regulatory network composed of *ZNF263*, *ZNF134*, *SP2*, *SP3*, and *VEZF1*, as the mostly shared TFs (Figure S2D). These results suggest a coordinated regulatory mechanism for DPGs in critical biological processes, such as cell cycle progression and chromatin remodeling. Notably, a higher number of co-expressed TFs is observed in tissue-specific co-expression of DPGs, whereas generally co-expressed DPGs do not show this pattern (Figure S2E). This result suggests that a greater number of TFs are required as cofactors for the complex, tissue-specific co-expression of DPGs.

### Conservation of Primate-associated DPGs suggests strong function-driven convergent evolution

Since DPGs show diverse lineage-associated patterns during evolutionary processes among vertebrates, we focus on four representative species within the Hominidae family, *Homo sapiens, Gorilla gorilla, Pan troglodytes*, and *Pongo abelii*. Using a straightforward sequence alignment to map out gain-and-loss events of lineage-associated and species-specific DPGs, we classified them into the four diversified types based on orthology and orientation. Despite the fact that DPGs of the four species are mostly shared at different degrees, there is still a set of orientation-specific DPGs (osDPGs), i.e., those that have lost their original orientations or exhibit altered spacing. From a gene-to-function point of view, lineage-associated DPGs are undoubtedly diverse in an absolute sense, especially between the pairs, but rather narrow in principle toward synchronization of core molecular mechanisms and cellular processes (Figure 6A–D; Table S7). This point is further highlighted by a genome sequence alignment of human and chimpanzee in multiple regions where functional enrichments are largely focused on replication (such as DNA synthesis) and transcriptional regulation (such as histone methylation) (Figure 6E). In addition, the 390 highly-conserved Hominidae-specific DPGs, at both gene and orientation levels, are enriched in functions of post-transcriptional (such as ncRNA processing) and translation processing (such as ribosome biogenesis) (Figure 6F). Furthermore, 12 human-associated DPGs appear to be best enriched in post-translational processing, such as protein-heterodimerization and chromatin construction (Figure 6G).

**Figure 6.**
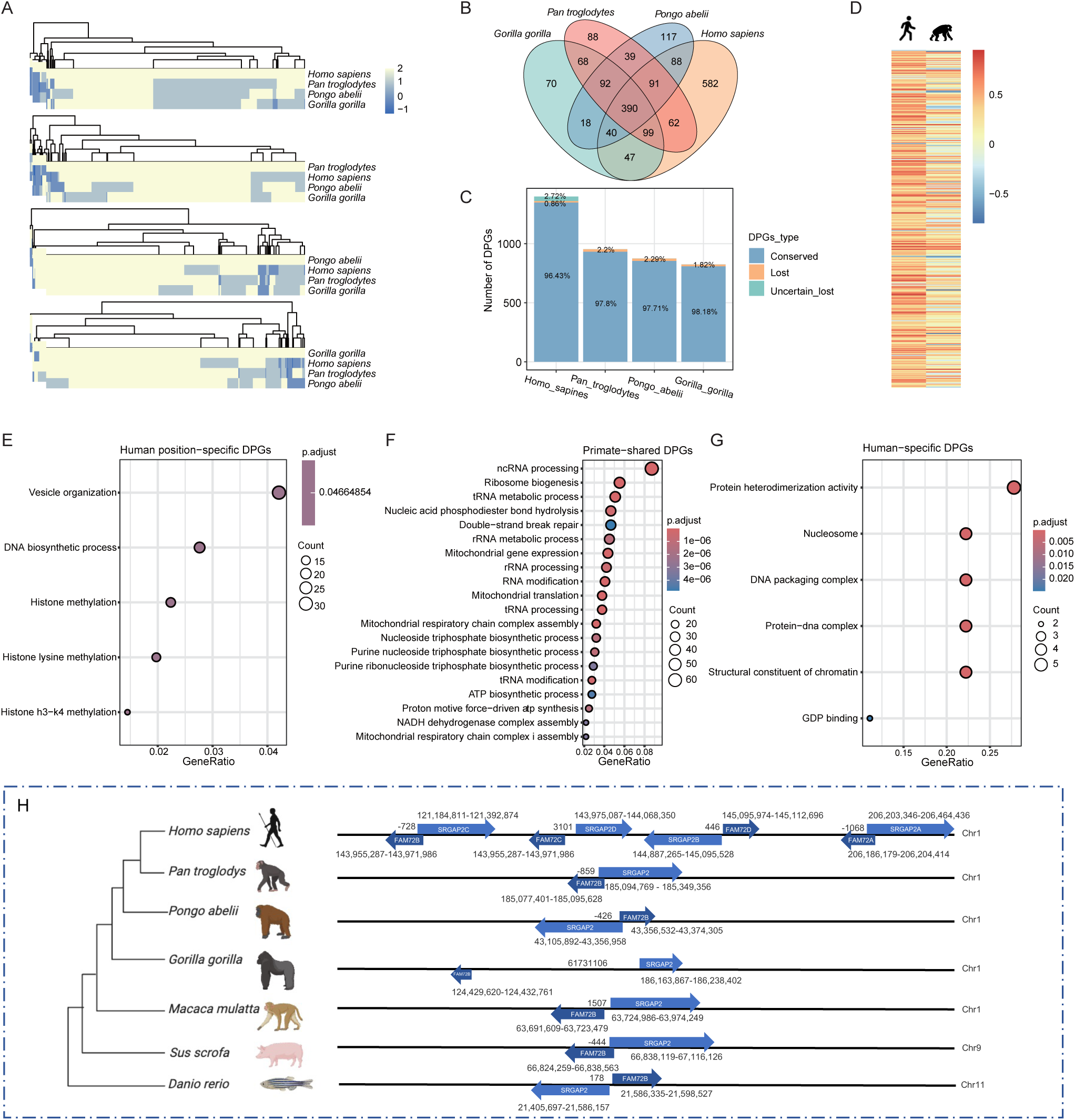
Conservation and specificity of the human and primate-associated DPGs. **A.** Heatmaps of DPGs showing the conservation across the four primate species. Each for a species shown in the first row. The darker color indicates lower conservation. The four values, ranging from 2 to −1, represent the four conservation forms as outlined in Figure 1, representing a conservation scale from high to low. The light blue block denotes the osDPGs in each species. **B.** A Venn diagram depicting shared and unique DPGs among the four species. **C.** A bar plot illustrating the counts of conserved and lost DPGs for each species. Conserved and Lost groups denote DPGs that are shared and lose the orientations at least one of the two counterparts of a DPG among the four primates, respectively. The human DPGs that are novel genes lacking annotation or repeatedly annotated at close positions, referred to as Uncertain-lost DPGs. The conserved DPGs in other three primates are similar, ∼ 98%. **D.** The co-expression correlation coefficient of human osDPGs in comparison with the pair of orthologs in chimpanzee. **E-G.** Functional enrichment of human osDPGs, primate-associated and human-associated DPGs. **H.** An Illustration of the human-associated DPGs, the *SRGAP2B-FAM72D* DPG family, and arrangements of their orthologous counterparts among other species.

Employing the macaque as a common ancestor, we further investigated the origin of interspecific sequence variations of the human osDPGs. The difference in osDPGs between human and chimpanzee originates from both deletions in human and insertions in chimpanzee within the inter-TSS sequence of DPG (Figure S3A). The insertion in chimpanzee displays an overrepresentation of repeat sequences of LINE and SINE elements (Figure S3B), with L1 and Alu being the most predominant types within each (Figure S3C and D).

Several human-specific DPGs are identified as playing a pivotal role in the development of the human species. Among these, the gene pair *SRGAP2-FAM72* is uniquely duplicated in human, leading to 4 copies in total resulting from two gene duplication events, whereas the DPG remains single in other vertebrate lineages (Figure 6H). Studies have shown that these DPGs are closely related to the development of nervous systems, such as the central cortex [37,38]. Also, human-associated DPGs play fundamental regulatory roles, such as *TUG1* as a notch-regulated gene [39].

### There is a diverse distribution of human-associated DPGs within and among human populations

T2T-CHM13 of European origin and the T2T-YAO genome of the Chinese population, have provided precise and comprehensive gene annotations for different populations. By comparing the genomes from different populations, we identified a group of distinct population-specific DPGs (psDPG). The results show that 55 and 357 DPGs are unique to YAO and CHM13, respectively (Figure S4A, Table S8). This outcome illustrates that DPGs not only exhibit unique patterns across different species but also display variations among populations. To assess the co-expression of psDPGs, we utilized RNA-seq data of healthy Chinese individuals obtained from the GSA database, accession number HRA004352 [40]. By mapping the RNA-seq data to each reference genome and calculating the Pearson correlation coefficient, we constructed a co-expression heatmap of these DPGs. The results reveal differences in co-expression among psDPGs (Figure S4B). These variations may represent potential mechanisms contributing to the distinctive traits observed in each population.

## Discussion

### Evolutionary significance of DPGs for future lineage-associated genome studies

Most, if not all, genes are non-randomly partitioned over chromosomes and chromatin territories; in general, the former are selected for replication and subsequent cell differentiation, and the latter are selected for transcription and cellular functions. In these regards, DPGs are an organizationally- and a functionally-diverse class of co-expressed genes, and their evolutionary studies are of the essence in understanding their mechanistic details and classifying them in lineage-associated and functionally-correlated ways [3,41–49]. In this study, we demonstrate these points of view by focusing our analyses on coarse-selected vertebrate lineages and in a human-centric design, as our previous investigations have been suggesting that evolutionary selectivity is biased between invertebrates (drosophila as examples in the case) and vertebrate lineages [3]. We further suggest that future DPG studies should pay special attention to systematic annotations of vertebrate DPGs as others have proposed for vertebrate biology and human disease studies [6,13]. In addition, DPG evolutions across eukaryotic lineages are both divergent and convergent in nature. In larger evolutionary scales, the highly conserved DPGs are mostly involved in highly conserved cell cycle or spatiotemporal-parameter-sensitive processes, through inventions of, by and large, direct binding or interaction for both relevant and irrelevant pathways and networks [4,10,50], whereas for closely related species or genre, many novel species- or lineage-associated DPGs appear to be generated for tissue- or organ-specific functions.

### Mechanistic implication of DPGs between species and populations

It is rather surprising that our comparative studies among closely related primates and between human populations reveal striking differences in DPGs, and such a result suggests that the in-and-out relationship created by forming and destroying DPGs is frequent in species evolution. Comparison between human and chimpanzee reveals that human psDPGs arise mostly biased sequence variations both in kind, such as deletion vs. insertion, and in number, such as more deletions in the human genome and more insertions in the chimpanzee genome during such a rather short evolutionary time scale. Moreover, The insertion of transposable elements (TEs) of Alu and L1 in chimpanzees contributed predominantly to the divergence between orthologs. Kanako O. Koyanagi et al have demonstrated that the emergence of DPGs involves exploiting pre-existing genes, showing consistency with our results [6]. Furthermore, we also identified that a portion of DPGs emerges specifically in certain species due to segmental duplications or gene duplications (Figure 6H). In addition, DPGs also show differences between human populations. Our results from a comparison between the two reference genomes, those of the Chinese and European populations, highlight the existence of variability and specificity among DPGs. These distinctive population-associated characteristics of DPGs may provide novel insights into how genomes should be analyzed and what targeted therapeutics should be developed for future medicine.

### Regulatory principles and functional novelties

The evolution with a limited repertoire of genes has resulted in complex life forms and multi-tracked holovivological systems, whose molecular details are believed to be involved in five distinct tracks: information, operation, homeostasis, compartmentalization, and plasticity [42,51]. On the informational track, DPGs can be mapped out systematically based on high-quality genome assemblies and such a mapping process can be extended in one dimension, eg., to construct databases for DPGs of all species, and in the other dimension, eg., to map DPGs between protein-coding and non-protein-coding gene pairs. Although the ultimate goal of such studies is to divide all genes either in a cluster or regulated alone, efforts toward the final ones have to be stepwise. On the operational track, the regulatory mechanisms of DPGs at different cellular processes are to be compared to those of non-clustered genes. Based on our current study, we are able to draw four basic conclusions. First, there have not been novel mechanisms in regulating DPG expression other than better coordinated expression at either the same or uneven levels. Second, there have not been many unique nor novel regulatory elements and mechanisms within the inter-TSS sequence spaces other than the enrichment of certain trans-regulatory TF classes and cis-regulatory sequence variations that are sensitive to sequence lengths, which are measurable based on insertion and deletion events according to to to population genetic conception. Third, the frequency distribution of indels within the inter-TSS sequences also shows a strong association with nucleosome phasing or space occupation, as shown in our previous work [52], and the phenomenon supports the two previous notions. Fourth, the length variability of the inter-TSS distances among the three types of DPGs confirms that it is not only selected but also has a tendency to maintain an optimal length for the best fit of their trans-regulatory elements. Interestingly, a similar trend was observed in small introns with optimal lengths, as highlighted in our previous work [53–57]. Our final notion is to emphasize that our study indicates the necessity for further investigation of DPGs on the other three holovivological tracks where their functional relevances to cell cycle timing and brain association are to be further inspected.

## Conclusion

Our discoveries underscore the essential role of DPGs in the human genome, presenting new avenues for understanding their contributions to genetic regulation and evolutionary biology. The observed patterns of conservation and selection suggest that DPGs are not only structural elements but also pivotal in maintaining genomic integrity and functionality. Future research should continue to explore the regulatory mechanisms and evolutionary trajectories of DPGs, providing deeper and more detailed insights into their biological significance and potential applications in genomics and medicine.

## Materials and methods

### Species selection and data collection

In order to conduct a conservation analysis of human DPGs in vertebrates, an additional 45 species were selected in a range of different evolutionary scales, including fish, amphibians, reptiles, mammals, primates, and birds. This was done following a quality screening with contig N50 values greater than 15k (Table S2). All species were from the Ensembl database (version 108) [58], including DNA sequences, protein sequences, and gff3 files for further analysis.

### Epigenetic data collection

The epigenetic data used in this study were sourced from the ENCODE project [59], accessed via the UCSC Table Browser [60] and the FTP site. The datasets for H3K4me3 and H3K27ac are wgEncodeRegMarkH3k4me3 and wgEncodeRegMarkH3k27ac, respectively, both obtained from UCSC. Six non-disease cell lines were retained from the two data sets, including the lymphoblastoid cell line (Gm12878), H1 human embryonic stem cell (H1-hESC), Skeletal Muscle Myoblast (Hsmm), human umbilical vein endothelial cells (HUVECs), normal human epidermal keratinocytes (NHEK), and normal human lung fibroblasts (NHLF). The signal values were calculated and averaged across the six cell types. The DNase dataset, wgEncodeRegDnaseClustered, was also downloaded from UCSC. RNAPII data were obtained from the ENCODE portal [61]. Three cell types were selected for analysis: GM12878, H1-hESC, and HUVECs. The accession numbers of the three datasets are ENCFF322DRU, ENCFF942TZX, and ENCFF696JRV.

### Orthologous genes identification

The orthologous gene relationships between species were identified by OrthoFinder [62] (version 2.5.4) with default parameters. Protein sequences from all species were used as input, and for genes with multiple transcripts, the longest transcript was retained. Orthologous genes provided by the Ensembl database were included as a complement in the comparative analysis of invertebrates and Hominidae species, and the data was retrieved via Ensembl Biomart [63]. In regard to the discrepancy between the two orthologous datasets, each is preserved. The combination method and data format are in reference to the HGD database [64].

### Identification and comparison of DPGs

To identify DPGs, we screened the protein-coding genes in the gene annotation files of each species. We found gene pairs that met the requirements of opposite transcription directions in the same chromosome and a distance between the TSSs of less than 1 kb. To construct a cross-species conservation map of DPGs, we searched for the corresponding orthologous genes in other species. We classified all DPGs into four types of conservation patterns: (1) highly conserved, referring to DPGs that are conserved across the species being compared and have orthologous genes maintain the same orientation; (2) conserved, referring to DPGs that are conserved across the species but one of the paired genes lost their original orientations; (3) lost single, referring to DPGs that lost one of the paired genes on an orthologous counterpart; and (4) lost both, both of the DPG pairs are missing in the original location. The R package pheatmap [65] was used to plot the conservation values of all DPGs across species. Hierarchical clustering was performed based on a complete linkage method to cluster rows of species. The comparison between different human reference genomes is similar to cross-species mapping. All DPGs in different reference genomes are identified and mapped to each other based on gene symbols. Unmapped DPGs will be referred to as psDPGs for further co-expression comparisons.

### House-keeping genes in DPGs

The list of HK genes was downloaded from a previous study [28]. The content of HK genes in DPGs was determined by matching HK gene with DPGs. The bitr function of the R package ClusterProfiler [66] was used for ID conversion. DPGs containing at least one HK gene were defined as HK-DPGs.

### Function enrichment analysis of DPGs

Gene function enrichment analysis was conducted using the R package Clusterprofiler [66]. The Benjamini-Hochberg (BH) method was employed to adjust the p-values, with a significance threshold of P < 0.05 and q < 0.05. The Database for Annotation, Visualization and Integrated Discovery (DAVID) (v6.8) [67] (https://david.ncifcrf.gov/) was employed to perform the various categories of gene function enrichment analysis, including Functional Category (e.g. UP KEYWORDS), Gene Ontology (e.g. GOTERM BP DIRECT, GOTERM CC DIRECT, GOTERM MF DIRECT), Pathway (e.g., KEGG PATHWAY) for candidate gene lists under a current background *Homo sapiens* with a two-tailed corrected Fisher’s Exact test (P < 0.05).

### PPI and RNA-protein interactions

The PPI data was retrieved from the STRING [68] database using the medium-confidence filtering threshold. In Figure 2D, only the two main clusters are retained and the dispersed nodes are reduced. The combined interaction score between the genes was used to calculate the edge thickness. The data on RNA-protein interactions was obtained from the RNAInter database (version 4.0) [69], which was filtered with a high confidence threshold of score > 0.2. The networks were constructed using the Cytoscape software, version 3.10.0 [70].

### Co-expression analysis of DPGs

Gene expression profiling data were obtained from the GTEx portal [71]. Samples were quality controlled using criteria of RIN > 6. The number of samples was counted after filtering, and only tissues with a sample number > 20 were retained (Figure S1C). Transcrips per million (TPM) expression profiles of the samples in all tissues were obtained, which were then transformed into natural logarithms log(TPM+1). The expression value TPM of each gene in a specific tissue represents the median TPM value across all samples. The two genes in DPGs were computed as the Spearman correlation between all the samples in each tissue using the spearmanr function of the Scipy package (version 1.11.2) [72]. The resulting values were organized into a matrix format and plotted as a heatmap using the Pheatmap package in R.

A filtering process was applied to identify the number of co-expressed DPGs in each tissue. DPGs with a co-expression coefficient greater than 0.8 and a P value less than 0.05 were retained. The fold differences in expression of DPGs were calculated by dividing the expression of highly expressed genes by the expression of lower expressed genes in the highest co-expressed tissue. The criteria for identifying the generally co-expressed genes were a co-expression coefficient greater than 0.7 in more than 70% of the tissues.

The RNA-seq dataset GSE127898 [73] was used to examine co-expression variation of psDPGs between human and chimpanzee in Figure 6D using a similar process.

### Epigenetic characteristics analysis

Regulatory figures for each pair of DPGs were plotted by the R package ggplot2. Each dataset mentioned above was calculated according to the specific position for each DPG. The DNA methylation site was set value of 20 as amplification. The DNase value was reduced by a factor of 10 for better display. GC count was calculated by sliding windows, with a window size of 50 bp and a step of 3 bp. When identifying the regulatory regions, we determined the distribution based on the relative positions of the TSSs and the peak of H3K4me3. The peak positions are identified by using the find_peaks function from the scipy.signal module.

The classification of DPGs is based on the highest co-expression correlation coefficient observed among all tissues. This classification is divided into three groups: high, moderate, and low, corresponding to correlation coefficients > 0.9, 0.8 ∼ 0.9, and < 0.8, respectively. For each group, the average signal chart was calculated for the four signal types, including DNase, RNAPII, H3K4me3, and H3K27ac. For each category, the distance between the TSSs of two genes in DPGs was normalized and uniformly scaled to 1000 bp. The value at each position was averaged for all DPGs, resulting in a single value for each position.

### Identification of shared TFs among DPGs

The TF motif data originates from the HOCOMOCO database [74] (version 11) and contains 769 full human TF motifs. The data were downloaded in meme format from MEME [75] (version 5.5.4). The binding site analysis was performed by using the FIMO [76] command-line version from the MEME suite. The analysis is based on motif file inputs along with sequence files of regulatory regions, with the selected parameter —norc to analyze the TFBSs on the strand where the gene resides. The co-expression of shared TFs is analyzed by the co-expression analysis method employed for DPGs. A filtering criterion with a correlation coefficient greater than 0.8 was used to identify co-expressed TFs as shown in Figure S2A.

### Identifying sequence changes of DPGs

Multiple sequence alignment was performed on the intergenic region of human osDPGs and the corresponding region between orthologous genes in human, chimpanzee, and macaque. The DNA sequence of the macaque was downloaded from the Ensembl database, the genome assembly is GCA_003339765.3. As for the sequences in the chimpanzee genome that do not align with the human genome, they are compared with macaque genome. Regions shared by chimpanzee and macaque but absent in human are defined as human-lost regions, while regions missing in macaque are defined as chimpanzee insertions. RepeatMasker [77] is used to analyze the repeat sequences in these specific regions of the chimpanzee genome.

The indels within the inter-TSS sequence spacing are derived from the 1000 Genome Project described in [36]. We filtered the indels based on a frequency threshold of greater than or equal to 0.005 and a length change of greater than or equal to 2 base pairs. All DPGs were grouped based on the inter-TSS distance of DPGs, ranging from −1k bp to 1k bp in 50-bp bins.

### Statistical calculations

All data were analyzed using Python (version 3.11.4), with the Pandas and Numpy packages utilized for data analysis, computation, and preprocessing. The Scipy package (version 1.11.2) [72] was used to calculate differences between groups using the Fisher test for small groups and the Chi-squared test for large groups (n > 40), both with P value < 0.05 as the criterion. Differences in distributions were tested by using the Wilcoxon-Mann-Whitney U test in R version 4.3.1.

## Code availability

The code for DPG identification has been compiled into a lightweight toolkit, which is available at https://ngdc.cncb.ac.cn/biocode/tool/BT7623.

## CRediT author statement

**Guangya Duan:** Data curation, Methodology, Formal analysis, Visualization, Writing – original draft, Writing – review & editing. **Sisi Zhang:** Methodology, Resources. **Bixia Tang:** Methodology. **Jingfa Xiao:** Methodology, Resources, Writing – review & editing. **Zhang Zhang:** Methodology, Resources, Writing – review & editing. **Peng Cui:** Conceptualization, Methodology, Resources, Writing – original draft, Writing – review & editing, Supervision. **Jun Yu:** Conceptualization, Methodology, Resources, Writing – original draft, Writing – review & editing, Supervision. **Wenming Zhao:** Conceptualization, Methodology, Resources, Writing – original draft, Writing – review & editing, Supervision. All authors read and approved the final manuscript.

## Competing interests

The authors have declared no competing interests.

## Acknowledgments

This work was supported by National Key R&D Program of China (Grant No. 2023YFC2605700 to WZ); Strategic Priority Research Program of Chinese Academy of Sciences (Grant No. XDB38050300 to WZ).

## Supplementary materials

**Figure S1** Co-expression patterns of vcDPGs **A.** A heatmap of co-expressed vcDPGs. DPG and expressed tissue are shown in rows and columns, respectively. The P values of the Spearman correlation coefficient are denoted as follows: *P < 0.05, **P < 0.01, ***P < 0.001. **B.** GO enrichment of generally co-expressed DPGs. **C.** Number of samples of each tissue after quality control.

**Figure S2** Vertebrate-conserved DPGs with shared co-expressed TFs among tissues **A.** A heatmap shows the shared TFs among DPGs. The blue color denotes shared TFBSs at regulatory regions of DPGs, whereas the yellow color does not. TFs (rows) and DPGs (columns) are grouped based on two-way hierarchical clustering. **B-C.** Functional enrichment of the two groups of most shared TF clusters. **D.** A network depicts co-expressed DPGs and shared TFs. Green triangles and yellow circles represent DPGs and TFs, respectively. Only the top three co-expressed TFs are shown. **E.** A heatmap shows co-expressed TFs shared by vcDPGs across various tissues, with a filtering criterion of co-expression value of Spearman correlation coefficient > 0.8 and P value < 0.05. Stars and colors indicate co-expression of DPGs and the number of expression-shared TFs, respectively.

**Figure S3** Comparison of inter-TSS sequence between human osDPGs and corresponding regions in chimpanzees **A.** The human osDPGs as compared to those of chimpanzee based on alignment to ancestor sequences referenced of the macaque genome. **B.** A pie chart illustrates the types of repetitive sequences in the region between orthologous counterparts of human osDPGs in chimpanzee. **C-D.** Types of LINE and SINE insertions between orthologous genes of human osDPGs in chimpanzee.

**Figure S4** A population-based analysis of different reference genomes **A.** A Venn diagram shows DPGs shared or unique to each human reference genome. Both haplotypes of the YAO genome are included. **B.** A heatmap illustrates the co-expression of all psDPGs. Rows represent DPGs, and the annotation bar denote their population specificities.

**Table S1** Lists of all protein-coding DPGs and vcDPGs involved in this study

**Table S2** A list of vertebrate species involved in this study

**Table S3** A list of human DPGs conserved in selected invertebrate species

**Table S4** Lists of DPGs involved in protein-protein, protein-DNA, and protein-ion interactions

**Table S5** A list of DPGs that are commonly co-expressed among different tissues

**Table S6** All tissues and corresponding abbreviations involved in this study

**Table S7** Lists of human osDPGs in types and unique human DPGs

**Table S8** DPGs identified in the three reference genomes

