## Supplementary figures and images for "Lineage-associated Human Divergently-Paired Genes (DPGs) Exhibit Regulatory Characteristics and Evolutionary Trends"

### Supplementary Figure 1

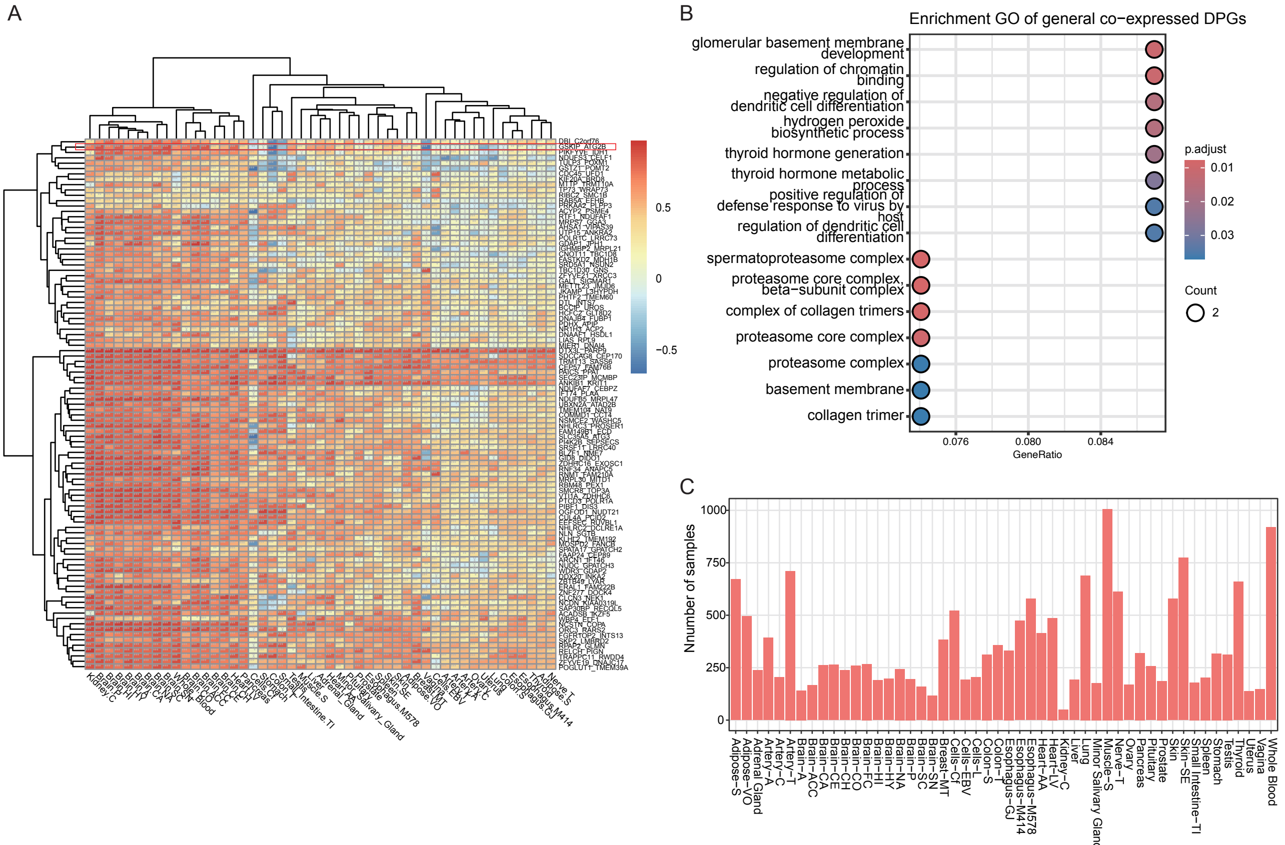

### Supplementary Figure 2

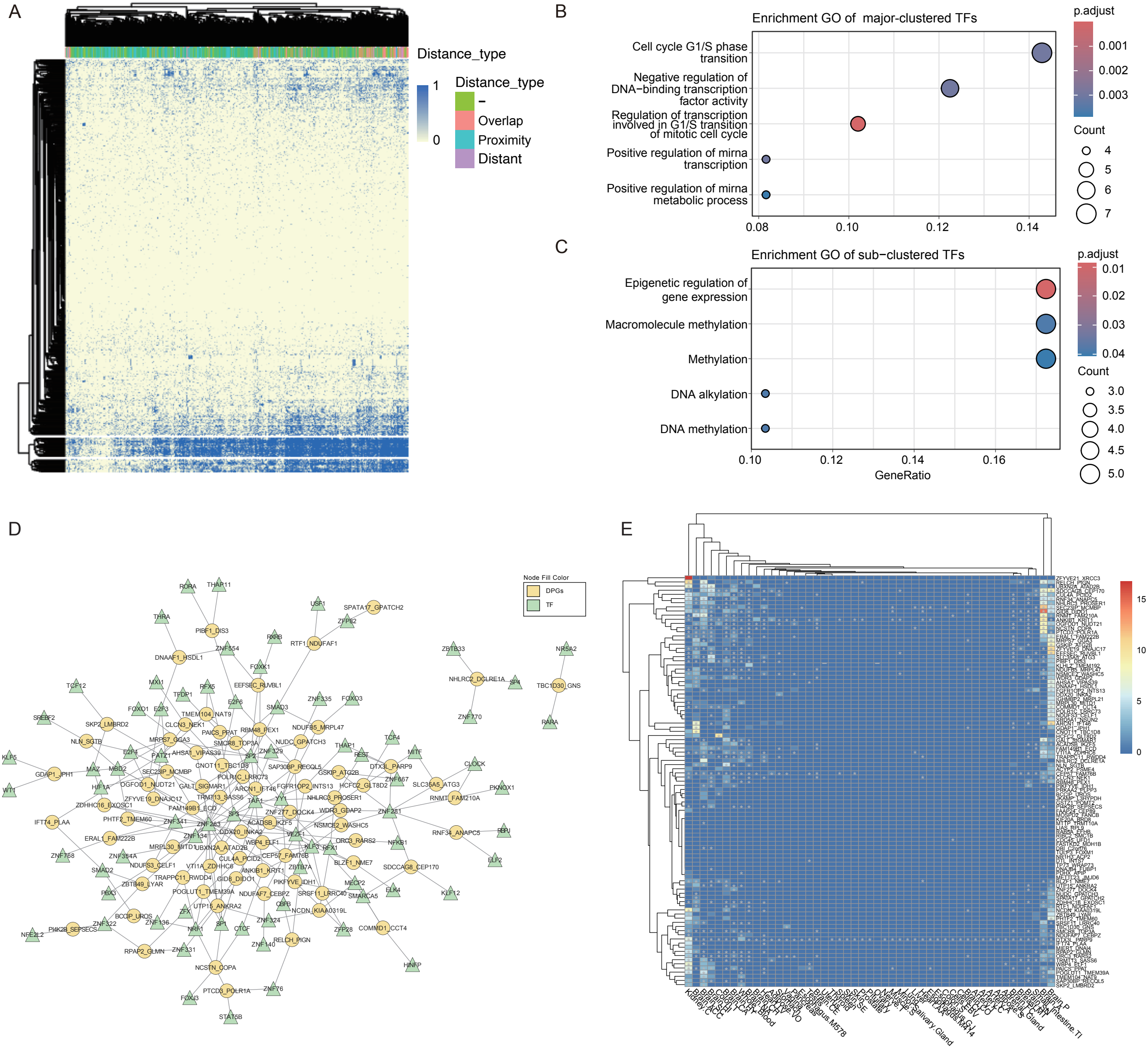

### Supplementary Figure 3

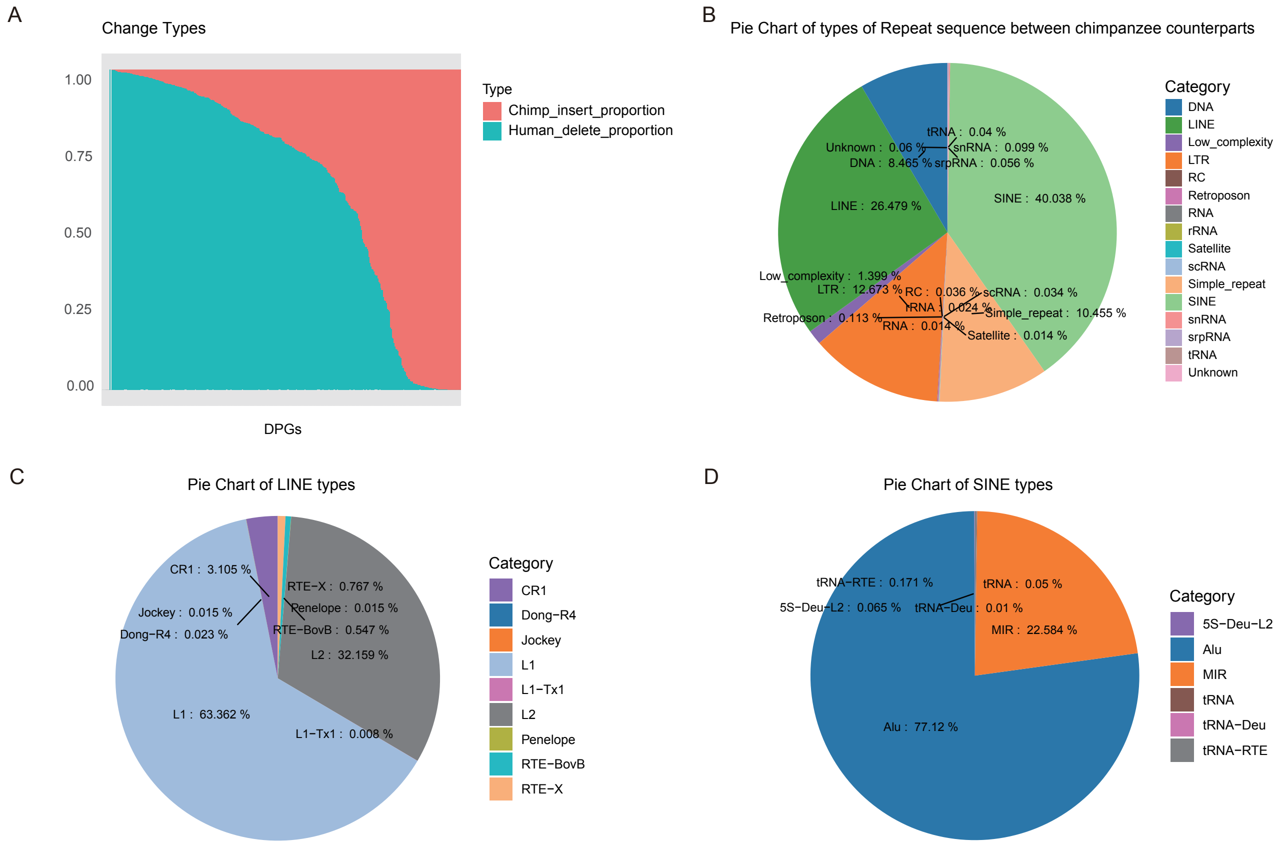

### Supplementary Figure 4

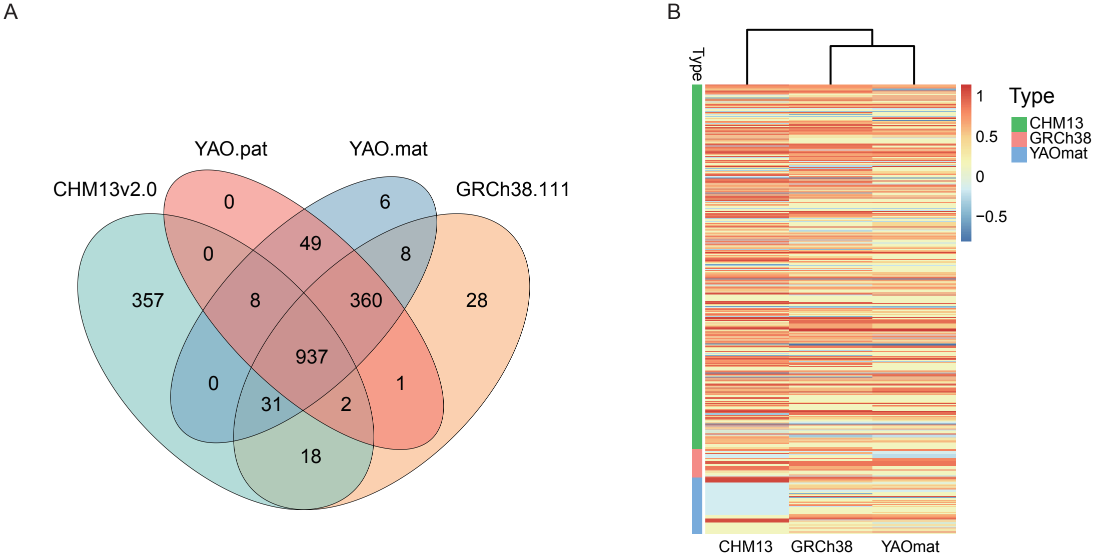
